# Early onset motor defects and electrographic seizures in a mouse model of the most common mutation in EEF1A2 related neurodevelopmental disorder, E122K

**DOI:** 10.1101/2023.09.07.556644

**Authors:** Grant F. Marshall, Melissa Fasol, Faith C.J. Davies, Matthew Le Seelleur, Alejandra Fernandez Alvarez, Cavan Bennett-Ness, Alfredo Gonzalez-Sulser, Catherine M. Abbott

## Abstract

*De novo* heterozygous missense mutations in *EEF1A2*, encoding neuromuscular translation-elongation factor eEF1A2, are associated with developmental and epileptic encephalopathies. We used CRISPR/ Cas9 to recapitulate the most common mutation, E122K, in mice. Although E122K/+ mice were not observed to have convulsive seizures, they exhibit frequent electrographic seizures and EEG abnormalities, transient early motor delays and growth defects. Both E122K homozygotes and *Eef1a2*-null mice develop progressive motor abnormalities phenotypes, with homozygotes reaching humane endpoints by P31. Surprisingly, E122K homozygotes did not exhibit the progressive spinal neurodegeneration which drives the null phenotype. The E122K protein is relatively stable in neurons yet highly unstable in skeletal myocytes, suggesting that the E122K/E122K phenotype is instead driven by loss-of-function in muscle. Importantly, E122K homozygotes developed abnormalities far earlier than nulls, suggesting a toxic gain-of-function. This novel mouse model represents the first animal model of an *EEF1A2* missense mutation with face-valid phenotypes and has provided mechanistic insights needed to inform rational treatment design.

## Introduction

Developmental and epileptic encephalopathies (DEE) are rare neurodevelopmental disorders (NDD) resulting from mutations in any one of over 200 genes (Helbig and Ellis, 2020), including *EEF1A2*. DEEs are characterised by early onset seizures with developmental delay or regression. In addition to the debilitating seizure burden, they are associated with other challenges including intellectual disability and behavioural problems, taking a substantial toll on those affected and their families (Gallop et al., 2021; Pressler and Lagae, 2020). Only a limited number of anti-seizure drugs are licensed for use in infants, and, while the rate varies by syndrome (McTague and Cross, 2013), many patients are refractory to treatment (Pressler and Lagae, 2020). There is a pressing need for new models of these disorders which show both construct and face validity in order to fulfil ambitions for precision medicine for severe, early onset epilepsy (Knowles et al., 2022; Marshall et al., 2021).

D*e novo* heterozygous missense mutations in *EEF1A2* resulting in DEE and NDD were first identified in 2012 using trio-based exome sequencing (de Ligt et al., 2012; Veeramah et al., 2013). Around 200 affected individuals have now been described in the literature or in clinical databases, with over 50 causative missense mutations found (Supplementary Figure 1). *De novo EEF1A2* mutations have now been estimated to occur with an annual incidence of 2.92 per 100,000 live births (López-Rivera et al., 2020), although the prevalence has not been estimated. The majority of cases have been identified in children, due to growing use of whole exome sequencing in paediatric populations presenting with early onset epilepsies and neurodevelopmental deficits. However, given that genetic screening of adults with NDDs is not yet widespread (Zacher et al., 2021), a substantial cohort of undiagnosed adults with *EEF1A2* missense mutations is likely to exist.

Most individuals with heterozygous missense mutations in *EEF1A2* experience intractable seizures that often begin in the first year of life, are minimally or completely non-verbal, and have moderate to severe intellectual disability (Carvill et al., 2020; Lam et al., 2016; Nakajima et al., 2015). Diverse seizure types are seen, including infantile spasms, nodding spasms, typical and atypical absences, myoclonic seizures, and focal or generalised tonic-clonic seizures (Carvill et al., 2020; de Ligt et al., 2012; Lam et al., 2016; Wang et al., 2020). Microcephaly and structural brain abnormalities are also seen in some patients, as are challenging behaviours and autism. Around half of patients are unable to walk and most of those with reduced mobility also present with movement disorders including ataxia, chorea, and dystonia. While several mutations have recurred in multiple cases, many have been found in only one individual (see Supplementary Figure 1). This makes genotype-phenotype comparisons difficult, but certain mutations are associated with more severe disease. Recent reports also suggest that some will develop a neurodegenerative course with poor long-term outcomes (Carvill et al., 2020). There is thus a real need for precisely targeted treatments in addition to standard anti-seizure medication.

The *EEF1A2* gene encodes neuromuscular translation elongation factor eEF1A2. The canonical role of eEF1A proteins is the GTP-dependent delivery of aminoacylated tRNAs to the A-site of the ribosome during protein synthesis, functioning as part of the elongation factor 1 complex in conjunction with the GTP exchange factor complex eEF1B (Li et al., 2013). All vertebrates possess two independently encoded eEF1A isoforms (eEF1A1 and eEF1A2) which, despite being 92% identical at the amino acid level (Soares et al., 2009) and performing almost equivalently in translational assays (Kahns et al., 1998; Timchenko et al., 2013), have distinct spatiotemporal expression patterns. During early embryonic development, eEF1A1 is expressed ubiquitously and seemingly exclusively, with eEF1A2 later emerging alongside eEF1A1 in embryonic neurons and muscle (Davies et al., 2023). Postnatally, a conserved isoform switch occurs in which eEF1A2 completely replaces eEF1A1 in cardiac and skeletal muscle (Chambers et al., 1998; Lee et al., 1992, 1993; Pan et al., 2004) and partially replaces eEF1A1 in neurons; neuronal somata switch entirely to eEF1A2, with eEF1A1 thereafter restricted to axons (Davies et al., 2023). The eEF1A isoform switch is typically complete by around postnatal day 21 (P21) in rodents (Chambers et al., 1998; Pan et al., 2004). The reason for this conserved tissue-specific switch between almost identical proteins remains unknown, but is hypothesised to relate to the differing non-canonical functional profiles of eEF1A1 and eEF1A2 (Mendoza et al., 2021; Mills and Gago, 2021).

A major consideration for *EEF1A2* missense mutations is to discover whether they broadly result in loss-of-function (LOF), gain-of-function (GOF), or both, as this would direct any therapeutic approach. While over 50 causative missense mutations in *EEF1A2* have been described, no clear LOF variants (small deletions or frameshifts) have been reported, which is a highly unusual mutational profile suggestive of GOF (McLachlan et al., 2019). The missense mutations are scattered throughout the gene and are found in every coding exon (Supplementary Figure 1). Although there is some enrichment in the GTP hydrolysis and tRNA binding domains, there is no strict clustering in particular domains from which we might infer the likely functional impacts. Disease models have revealed important functional insights about select *EEF1A2* mutations, however, the degree of LOF/ GOF and the molecular mechanisms involved are only beginning to be established. For example, studies in yeast have concluded that the main mechanism for a number of variants is simple LOF (Carvill et al., 2020). Functional studies in mammalian systems are non-trivial, as all transformed, immortalised, iPSC-derived, and primary cell cultures so far studied express eEF1A1, either exclusively or alongside eEF1A2, confounding genotype-phenotype correlations and potentially obscuring any LOF/ GOF (McLachlan et al., 2019). A successful strategy we have previously employed is comparative phenotyping of mice carrying missense and null mutations in *Eef1a2* on the same genetic background (Davies et al., 2017, 2020). Comparison of weight gain profiles and neurological scores in mice homozygous for the D252H missense mutation and mice homozygous for null mutations revealed that mice carrying D252H were consistently more severely affected than nulls, suggesting a GOF. Meanwhile, mass spectrometry analysis revealed that D252H also prevents eEF1A2 from binding its cognate guanine exchange factor eEF1B, which is likely to cause a LOF (Davies et al., 2020). While mouse models have thus revealed important functional insights, a key issue with those described so far is the lack of face valid phenotypes (seizures, electroencephalographic abnormalities, motor abnormalities, learning and memory deficits, autism-relevant behaviours) with which to assess the efficacy of experimental therapies, particularly in the clinically relevant heterozygous missense mice. Mouse and human eEF1A2 only differ by a single amino acid, meaning that any of the causative missense mutations described so far can be straightforwardly modelled in mice, and recapitulating more clinically severe mutations may be more likely to produce models with face valid phenotypes. The E122K mutation (c.364G > A, p.Glu122Lys) has been described in at least 11 individuals in the literature, making it the most commonly identified mutation. E122K presents with a consistently severe phenotype (published clinical findings summarised in Supplementary Table 1). All patients have early onset epilepsy, with onset typically in the first months of life and variable seizure control. There is strong evidence of epileptic encephalopathy in three cases, in which developmental arrest/ regression occurred as the epilepsy progressed. All individuals have moderate to severe intellectual disability. Affected children typically have movement disorders, though many can walk. Most are non-verbal and several are reported to show self-injurious behaviour, sleep abnormalities and autistic behaviours.

We used CRISPR/ Cas9 to generate a novel mouse model of the E122K mutation and carried out detailed phenotyping of both homozygotes and heterozygotes, alongside mice carrying null mutations on the same genetic background. Here we describe early motor delays, growth defects and electrographic seizures along with electroencephalogram (EEG) abnormalities in this model, and present evidence that E122K exerts a toxic GOF/ dominant-negative effect.

## Materials & Methods

### Animal methods

All mouse work was carried out in accordance with the Animals (Scientific Procedures) Act 1986 using protocols approved by the local ethics committee of the University of Edinburgh. All mice used in this study were maintained on the C57BL/6JCrl genetic background and maintained as inbred colonies. All mice were housed in the Western General Hospital Biological Research Facility (BRF) except the mice used for EEG, which were housed in University of Edinburgh Centre for Discovery Brain Sciences facilities. Mice in the BRF were kept in Blue-Line 1285L IVC cages (Techniplast) with wood chips and tissue paper for bedding. Cages were enriched with a clear polycarbonate tube (Datesand) and a wooden chew stick. All mice were kept on a standard 12-hour light 12-hour dark cycle (light on at 7 a.m. and off at 7 p.m.) at 18-24°C and 45-65% humidity and had *ad libitum* access to food and water. Mouse pups were ear-notched between P12 and P14 for identification/ genotyping and weaned after P21. Post-weaning animals were housed in single-sex mixed genotype cages containing 2-5 mice. Mice in the E122K line were typically supplied with mash diet at weaning to support the mutants.

All phenotyping and analysis was performed by experimenters blind to genotype. Male and female animals were used in all experiments. Mice were randomly assigned to each sex and genotype by Mendelian inheritance. Behavioural and motor tests were performed on animals in single-sex mixed genotype cages, with animals typically tested in the order of their colony number (randomly assigned at ear notching).

Throughout this paper, mice heterozygous for the E122K mutation in Eef1a2 (Eef1a2^+/E122K^) are referred to as E122K/+, while mice homozygous for the E122K mutation (Eef1a2^E122K/E122K^) are referred to as E122K/E122K. A second line of mice carrying the Del22Ex3 null mutation in *Eef1a2* (Davies et al., 2020) was studied alongside mice from the E122K line. Mice heterozygous for the Del22Ex3 mutation (Eef1a2^+/-^) are referred to as +/-, while mice homozygous for the Del22Ex3 null mutation in Eef1a2 (Eef1a2^-/-^) are referred to as -/-.

### Generation of Transgenic Mice

Guide crRNA 5’-CGCCTCAAACTCGCCCACAC-3’ was selected by entering the genomic sequence of *Mus musculus Eef1a2* (assembly GRCm38.p6) into the CHOPCHOP tool (Labun et al., 2019). We designed two 200-nucleotide single-stranded oligodeoxynucleotide (ssODN) repair templates using the genomic sequence of *Eef1a2*. The first repair template contained three single nucleotide substitutions resulting in E122K missense, silently abolished the protospacer-adjacent motif (PAM) to prevent repeated Cas9 mediated cleavage (Paquet et al., 2016), and silently incorporated a restriction site for the endonuclease PstI (facilitating convenient and low-cost genotyping of transgenic animals by polymerase chain reaction (PCR) and subsequent restriction digest (Abbott, 1993)). As biallelic mutations in *Eef1a2* cause postnatal lethality in mice (Chambers et al., 1998; Davies, 2017; Davies et al., 2017; Shultz et al., 1982), we designed and co-delivered a second repair template encoding only the silent PAM mutation in order to reduce the likelihood of generating such genotypes in the F0 generation (Davies et al., 2020). Repair templates were designed to be non-complementary to the crRNA (DiCarlo et al., 2013), and the introduction of rare codons (Zhao et al., 2021) was avoided.

#### E122K ssODN repair template (5’-3’)

5’∼TCTTGTTGACACCCACAATGAGCTGCTTCACACCCAGAGTGTAGGCCAGGAGTGCGTGTTCCCGGGTTTGCCCGTTCTTGGAGATGCCCGCCTCAAACTTGCCCACACCTGCAGCCACGATCAGCACTGCGCAGTCCGCCTAGCCAACAGGTCAGACACAGTGAGTCCCCACCCGGCCCTGCCTTCGACCTGGCCCTGCC∼3’

#### Second ssODN repair template, with only silent PAM mutation (5’-3’)

TCTTGTTGACACCCACAATGAGCTGCTTCACACCCAGAGTGTAGGCCAGGAGTGCGTGTTCCCGGGTTTGCCCGTTCTTGGAGATGCCCGCCTCAAACTCGCCCACACCTGCTGCCACGATCAGCACTGCGCAGTCCGCCTAGCCAACAGGTCAGACACAGTGAGTCCCCACCCGGCCCTGCCTTCGACCTGGCCCTGCC

E122K founder mice were generated by perinuclear microinjection of crRNA annealed to tracrRNA (Integrated DNA Technologies, 20 ng/µL) + Cas9 nuclease (Integrated DNA Technologies #1081058, 17 ng/µL) ribonucleoprotein complexes along with ssODN repair templates (Integrated DNA Technologies, each 75 ng/µL) into fertilized C57BL/6JCrl oocytes (Quadros et al., 2016). Microinjections were performed by staff at the Evans Transgenic Unit (Western General Hospital, Edinburgh). After 24 hours, surviving embryos were implanted into pseudopregnant CD1-IGS female recipients.

### Genotyping

DNA was extracted from ear notches using the HotSHOT method (Truett et al., 2000). All Sanger sequencing was performed by staff at the IGMM Technical Services Sequencing Service using a 3130 or 3730 Genetic Analyser (Applied Biosystems).

F0 animals were genotyped by amplifying exon 4 of *Eef1a2* with primers mE122KTOPOF and mE122KTOPOR and a proofreading polymerase (Platinum SuperFi DNA Polymerase (Invitrogen #12351250)), using the following thermocycler program:

> <u>98°C for 1 min, 35x (98°C for 5 sec, 64°C for 10 sec, 72°C for 15 sec), 72°C for 2 min, 10°C hold</u>

Blunt-end topoisomerase-based cloning was performed using the PCR products (Zero Blunt TOPO PCR Cloning Kit (ThermoFisher Scientific #K2875J10)), allowing the various *Eef1a2* alleles from each founder to be individually Sanger sequenced.

Further on target analysis was performed by amplifying a 1025 bp region of *Eef1a2* from an E122K homozygote with primers mE122K1kbF and mE122K1kbR and a proofreading polymerase (Platinum SuperFi DNA Polymerase (Invitrogen #12351250)), using the following thermocycler program:

> <u>98°C for 1 min, 35x (98°C for 5 sec, 54°C for 10 sec, 72°C for 15 sec), 72°C for 2 min, 10°C hold</u>

PCR products were verified by electrophoresis then Sanger sequenced using the amplification primers.

To perform off-target analysis, the ten loci in the mouse genome with the highest probability of cleavage by *S. pyogenes* Cas9 using the selected guide crRNA were identified using CRISPOR (Concordet and Haeussler, 2018). PCR primers were designed for these 10 loci (Table 1), which were amplified using a proofreading polymerase (Platinum Superfi II DNA Polymerase (Invitrogen #12361010)) using the following thermocycler program:

> <u>98°C for 1 min, 10x (98°C for 5 sec, 65°C for 10 sec (-0.5°C/cycle), 72°C for 15 sec), 33x (98°C for 5 sec, 60°C for 10 sec, 72°C for 15 sec), 72°C for 2 min, 10°C hold</u>

**Table 1.**
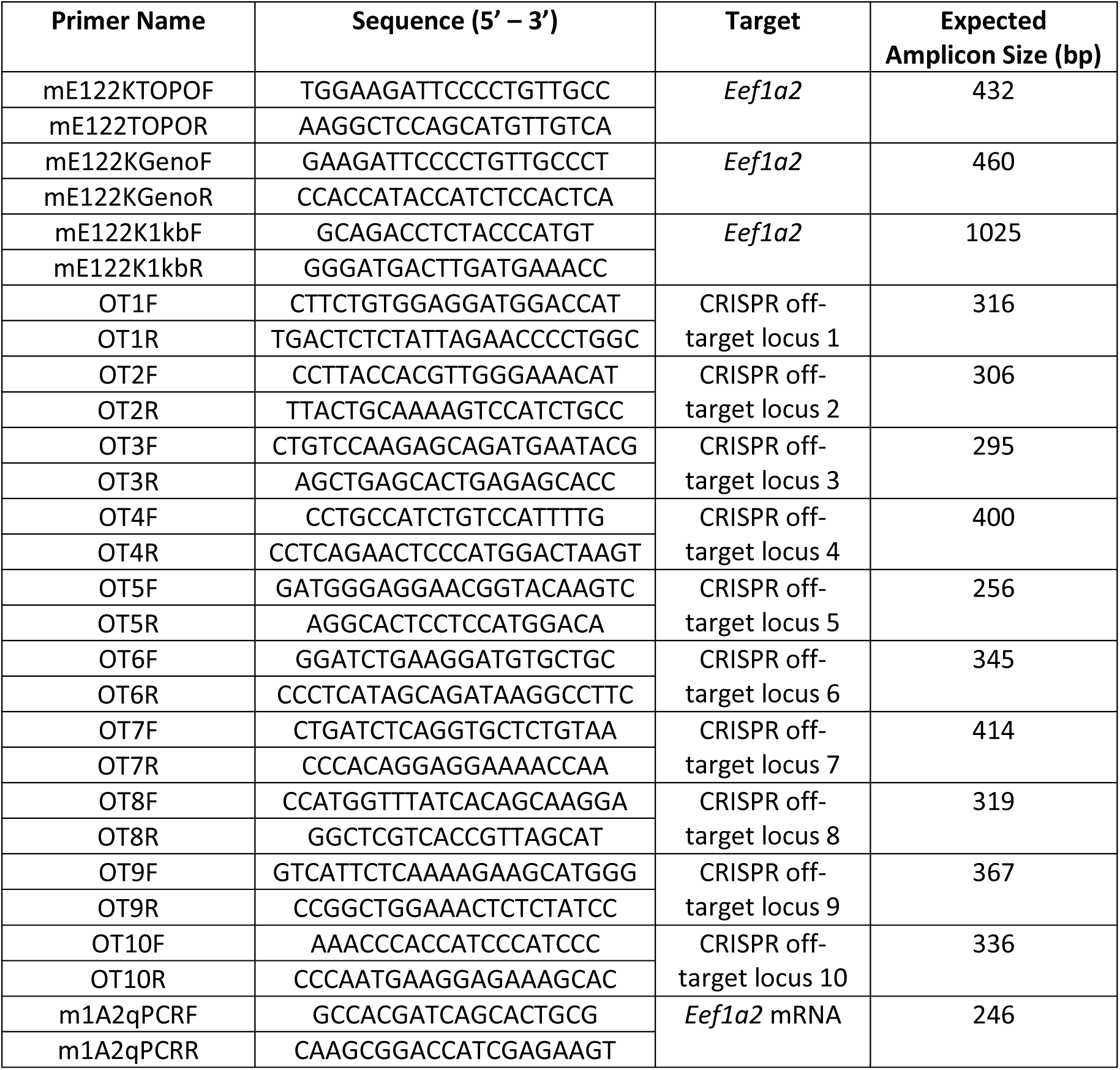
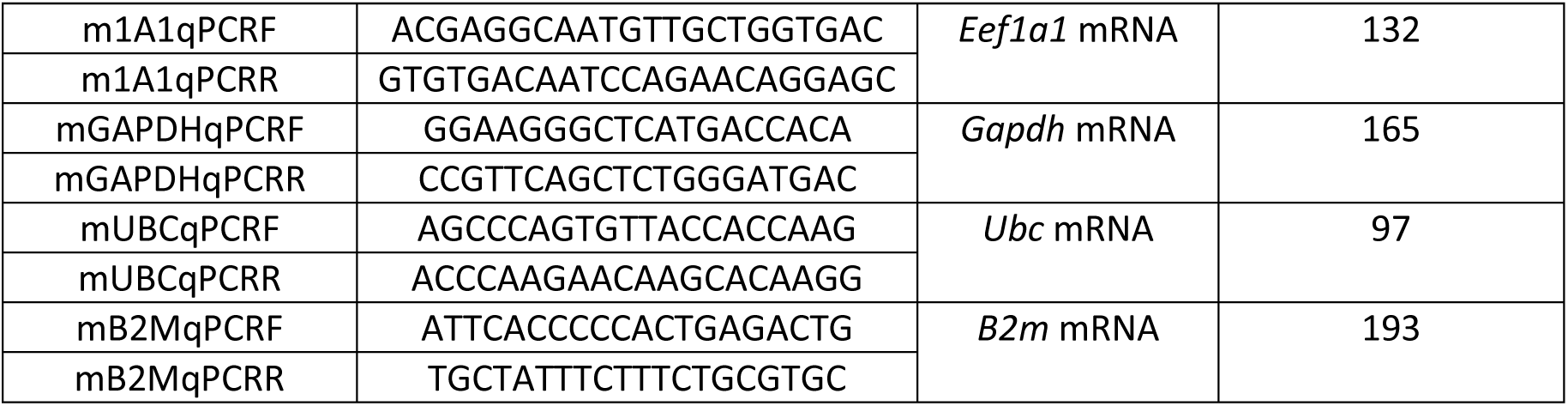
list of primers used.

PCR products were verified by gel electrophoresis then Sanger sequenced using the amplification primers.

Mice in the established line were genotyped by amplifying a region of *Eef1a2* centred on codon 122 with primers mE122KGenoF and mE122KGenoR and Taq DNA Polymerase (Life Technologies #10342020), using the following thermocycler program:

> <u>95°C for 2 min, 10x (95°C for 15 sec, 65°C (-1°C/ cycle) for 20 sec, 72°C for 50 sec), 25x (95°C for 15 sec, 55°C for 20 sec, 72°C for 50 sec), 72°C for 5 min, 10°C hold.</u>

PCR products were then digested using FastDigest PstI (ThermoFisher Scientific #FD01614) for 2-5 minutes at 37°C then analysed using gel electrophoresis. Genotyping by PstI digest is illustrated in Supplementary Figure 2D.

### List of Primers Used

### RNA Methods

Dissected tissues used for expression analysis were snap frozen on dry ice then stored at -70°C until use. RNA was extracted from P24 brain tissue (frontal lobe) using the Direct-Zol Miniprep Plus kit (Zymo #R2070T) according to the manufacturer’s instructions. Additional off-column DNAse digestion was performed using the DNA-*free*™ DNA Removal Kit (Ambion #AM1906. RNA was converted to complementary DNA (cDNA) using the High-Capacity cDNA Reverse Transcription kit (Applied Biosystems #4368814) according to the manufacturer’s instructions. Primers for reverse-transcription PCR (RT-PCR) and quantitative RT-PCR (qRT-PCR) were supplied by Merck with standard desalting at 100 µM in TE buffer. Primers and cDNA samples were validated by RT-PCR with the primers given in Table 1 and Taq DNA Polymerase (Life Technologies #10342020), using the following thermocycler program:

> <u>95°C for 2 min, 30x (95°C for 10s, 60°C for 30s, 72°C for 30s), 72°C for 7 min, 10°C hold</u>

qRT-PCR reactions were performed using Brilliant II SYBR Green qPCR Master Mix (Agilent #600828) according to manufacturer instructions using a Light Cycler HT7900 (Roche). ∼2 ng of template cDNA was used per qRT-PCR reaction, and the final concentration each primer was 300 nM. All reactions were run in triplicate, alongside -RT and water controls for each primer pair. In addition, standard curves were generated for each primer pair using a dilution series of wild-type cDNA (diluted 1 in 5, 50, 500, 5000 and 50000). qRT-PCR reactions were run on the following lightcycler program:

> <u>50°C for 2 min, 95°C for 10 min, 40x (95°C for 30s, 60°C for 1 min), 95°C for 15s, 60°C for 15s</u>

qRT-PCR data was analysed using the HT7900 SDS software (v2.4.2). Absolute quantities of the target gene in each sample were extrapolated from the standard curves using the threshold cycle (Ct) values, and then normalised to the geometric mean of the reference genes *Gapdh, Ubc* and *B2m* (Vandesompele et al., 2002). Melt curves were generated for each primer pair at the end of the thermocycler program.

### Protein Methods

For consistency, total protein was extracted from the parietal lobe of the brain, the whole heart, or a sample of hindlimb skeletal muscle containing *biceps femoris*, *vastus lateralis* and *gastrocnemius* muscle. Samples were lysed in a protein extraction solution (∼10 µL/ mg tissue) consisting of 0.32M sucrose with protease inhibitors (Merck #4693159001) and phosphatase inhibitors (Pierce # A32957) using a Precellys-24 lyser. Lysed samples were centrifuged at 10000 x *g* for 15 minutes at 4°C and the supernatants collected. 2x Laemmli loading buffer was added in a 1:1 ratio then proteins were denatured by heating to 100°C for 5 minutes on a heat block. 10% (v/v) of 1 M dithiothreitol was then added to each sample. For SDS-PAGE, samples were electrophoresed through 10% acrylamide gels in Mini Gel Tanks (Invitrogen) filled with 1x tris-glycine-SDS running buffer. 5 µL of Colour Prestained Protein Standard (New England Biolabs #P7719S) was included on each gel. Wet transfers onto Immobilon-FL PVDF Membrane (Merck # IPFL00005) were then performed using 1x NuPAGE transfer (Invitrogen #NP00061) buffer in Mini Gel Tank Blot Modules (Invitrogen). Total protein was stained using REVERT 700 total protein stain (LI-COR #926-11021) according to manufacturer instructions and visualised using a LI-COR Odyssey CLx Imaging System (169 µm resolution, low quality setting). Blots were de-stained then blocked for 1 hour at room temperature using INTERCEPT (TBS) Blocking Buffer (LI-COR #926-11021). Blots were incubated in primary antibodies diluted in INTERCEPT (TBS) Blocking Buffer overnight at 4°C. Primary antibodies were a custom-made rabbit-anti eEF1A2 (1:2000, see Supplementary Methods) or a custom-made sheep anti-eEF1A1 (1:1000) (Newbery et al., 2007). Blots were rinsed 3x for 10 minutes in TRIS buffered saline + 0.1% Tween20 (TBS-T). Secondary antibodies were applied in INTERCEPT T20 Antibody Diluent (LI-COR #927-65001) for 1 hour at room temperature in the dark. Secondary antibodies were IRDye 680LT Donkey anti-Rabbit IgG (1:20,000) (LI-COR #926-68023) or IRDye 800CW Donkey anti-Goat IgG (1:20,000) (LI-COR #926-32214). After a second set of washes, blots were imaged as described above and were analysed using Image Studio Lite (version 5.2). Band intensities was measured against background and normalised to a standardised section of total protein within each lane.

### Spinal Cord Histology

Spinal cords were isolated and immersion-fixed in 4% paraformaldehyde (in phosphate-buffered saline) overnight at 4°C then processed into paraffin blocks. 5 µm sections were taken from the cervical regions and mounted on glass microscope slides. Sections were stained with haematoxylin & eosin (H&E) using a standard protocol (Fischer et al., 2008) and coverslipped using a dibutylphthalate polystyrene xylene mounting medium. Brightfield images were acquired using a Hamamatsu C9600-02 NanoZoomer Digital Pathology microscope at 40x magnification using the default brightfield settings.

### Neuroscore Assessments

The “neuroscore” is a composite neurological function score originally designed for mouse models of spinocerebellar ataxia (Guyenet et al., 2010) but which has proven useful for phenotyping *Eef1a2* mutant mice (Davies et al., 2020). Briefly, hindlimb clasping, spinal kyphosis, ledge walking, and gait quality are individually scored between 0 and 3. These four scores are then summed to give a neuroscore ranging between 0 and 12, with higher scores representing more severe neurological phenotypes. Neuroscore criteria are detailed in the Supplementary Methods.

### Neonatal Motor Tests

Testing procedures for ambulation scoring, righting reflex tests and negative geotaxis tests were adapted from (Feather-Schussler and Ferguson, 2016). In brief, for ambulation scoring, the degree of gait development of P10 pups was scored on an ordinal scale between zero and 3, with higher scores representing more developed gaits. For righting reflex tests, P8 pups were placed in a supine position and the time to return to a prone position was measured. For negative geotaxis tests, P10 pups were placed facing down a 45° incline and the time taken to turn and face up was measured. Righting reflex and negative geotaxis measurements were performed 3x per pup with animals returned to the home cage for a minimum 2-minute break between tests.

### EEG/ EMG Electrode Implantation Surgery

Electrode implantation surgeries were performed after 6.5 weeks of age. Animals were anaesthetised with isoflurane and mounted on a stereotaxic frame (David Kopf Instruments, USA). Pairs of local field potential (LFP) electrodes (Teflon coated stainless steel, Ø = 50.8 μm. A-M Systems, USA), support screws and electrical ground screws (Yahata Neji, M1 Pan Head Stainless Steel Cross, RS Components, Northants, UK) were implanted at the coordinate locations given in Table 2.

**Table 2.** Stereotaxic coordinates for EEG recording electrodes and screws. All stereotaxic coordinates were calculated relative to bregma on the skull surface. The targeted hemisphere is indicated.

| Electrode/ Screw Location | Anteroposterior (mm) | Mediolateral (mm) | Dorsoventral (mm) |
| --- | --- | --- | --- |
| Motor Cortex (left) | 1.55 | -1.88 | Cortical surface |
| Somatosensory Cortex (right) | 1.3 | 2.0 | Cortical surface |
| Parietal Cortex (right) | -1.47 | 1.0 | Cortical surface |
| Medial Hippocampus (left) | -2.8 | -1.8 | 2.5 |
| Cerebellum (midline) | -6.0 | 0 | Cerebellar surface |
| Anterior support screws (bilateral) | 0 | ±3.5 | Epidural |
| Cerebellar ground screws (bilateral) | -6.0 | ±2.0 | Cerebellar surface |

Implanted electrodes were sealed in place using ultraviolet light-activated dental cement (3M Relyx Unicem 2 Automix, Henry Schein, UK). The electrodes were pinned to an electrode interface board (EIB-16. Neuralynx, USA) which was mounted on the top of the skull using dental cement. Finally, a stainless-steel wire was sutured onto the trapezius muscle for electromyogram (EMG) recording. Animals were allowed to recover for at least seven days before tethered recordings took place. Analgesia (Carprofen) was given perioperatively and then postoperatively as necessary.

### Tethered Polygraph and Video Recordings

Tethered EEG recordings took place between 7.5 and 12.5 weeks of age. Animals were continuously recorded during the light phase for 3.5-6 hours on at least two different days. Recordings took place inside (W x D x H) 50 x 50 x 40 cm perspex arenas, with wood chips as bedding. A small pot of water and some chow pellets were added to each arena. Up to four animals were recorded at once, with the animals unable to see one another and the experimenter blind to genotype. The electrode interface boards were tethered to RHD 16-channel recording headstages (Intantech, USA) wired to an acquisition board (OpenEphys, USA) via an electrical commutator (Adafruit, Italy). LFPs were recorded in the OpenEphys continuous format with a sampling frequency of 1 kHz (high pass filter >7500 Hz and low-pass filter < 2 Hz) and referenced to ground. Mice were simultaneously video recorded from above at ∼10 frames per second using a C270 HD Webcam (Logitech, USA).

### Analysis of EEG/ EMG recordings

EEG/ EMG analysis was performed in Python (V2.6.6) using custom code based on the MNE-Python package (V1.0.0) (Gramfort et al., 2013). First, continuous EEG/ EMG recordings were split into 5 second epochs. An experimenter blind to genotype classified epochs as wake, NREM or REM using the EEG and EMG waveforms using the following criteria based on EEG/ EMG characteristics. Wake was identified by the presence of desynchronized EEG and varying levels of EMG. NREM epochs displayed high-amplitude slow-wave (∼1–4 Hz) EEG activity accompanied by sleep spindles (∼12–17 Hz) and decreased EMG activity. REM was identified by sustained theta (∼5–10 Hz) and no EMG activity. The percentage of time in wake, NREM and REM, as well as the frequency of NREM/ REM bouts and the latency to REM were then calculated for each animal. NREM bouts were defined as any transition from waking or REM to NREM. REM bouts were defined as any transition from NREM to REM. Epochs with putative abnormalities (monospikes or polyspikes) were noted during vigilance state classification and were later revisited by a blinded experimenter for validation as normal, abnormal, or artefactual. Precise onset and offset times for validated polyspikes were determined and the average duration was quantified for each animal. For video analysis of cortical spike trains, polygraph and video recordings were synchronised using file metadata. Alignment was verified for each recording by studying sleep-to-wake transitions.

### Statistical Analysis

Statistical analyses were performed in Graphpad PRISM (Version 9), SPSS (Version 25) and R (Version 4.0.5). For all data that were statistically analysed, outlier removal was performed using the ROUT method (Motulsky and Brown, 2006), with a *Q* value of 1%. The normality of residuals was tested using the D’Agostino-Pearson omnibus (K2) test or Shapiro-Wilk test, as appropriate. The equality of variances/ homoscedasticity of data was also tested using Bartlett’s test of homogeneity of variances, the Brown-Forsythe test or Spearman’s rank correlation test, as appropriate. Appropriate parametric or non-parametric statistical tests were then performed. Parametric tests included one and two-way ANOVA and t-test (with or without Welch’s correction). Non-parametric tests included the Mann-Whitney U, Kruskall-Wallis and chi-square tests. Post-hoc testing (Dunn’s multiple comparisons test or Dunnett’s [T3] post-hoc test) and correction for multiple comparisons (Bonferroni’s correction or Holm-Šídák method) were performed as appropriate. Data transformation was performed where necessary to meet the assumptions of statistical tests. Individual animals were the statistical unit for all tests unless otherwise stated.

## Results

### Introducing the E122K mutation into the mouse *Eef1a2* gene

We used the clustered regularly interspaced short palindromic repeats (CRISPR) and CRISPR associated 9 (Cas9) system to generate the E122K mouse line. A schematic of the CRISPR design for E122K missense incorporation is given in Supplementary Figure 2. Four F0 mice carried E122K alleles, complete with silent PAM mutation and novel PstI site. Two of these F0 animals survived to breeding age and were mated with C57BL/6J stock, with one F0 animal transmitting the E122K allele to a single F1 female which was used to establish the E122K breeding line. DNA from this F1 heterozygote was used to screen for off-target Cas9-induced mutations, with none detected. DNA from an E122K homozygote was later used for on-target analysis, confirming that there were no unintended edits to *Eef1a2* in exon 4 or its flanking introns (Supplementary Figure 3).

### The E122K mutation reduces protein stability in a tissue dependent manner

All expression analysis was performed using tissue samples collected at P24, when the eEF1A isoform switch is typically complete in mice (Chambers et al., 1998; Pan et al., 2004). We found no difference in relative *Eef1a1* transcript levels in brains of mice with different genotypes (Figure 1A), indicating that the E122K mutation causes neither a delay in the *Eef1a* isoform switch or compensatory *Eef1a1* upregulation. Relative *Eef1a2* transcript levels were normal in E122K/+ mice but significantly increased in E122K/E122K mice (Figure 1B). There was therefore no indication that E122K and the associated silent mutations reduce transcript stability, although there is upregulation of *Eef1a2* in homozygote brains.

**Figure 1.**
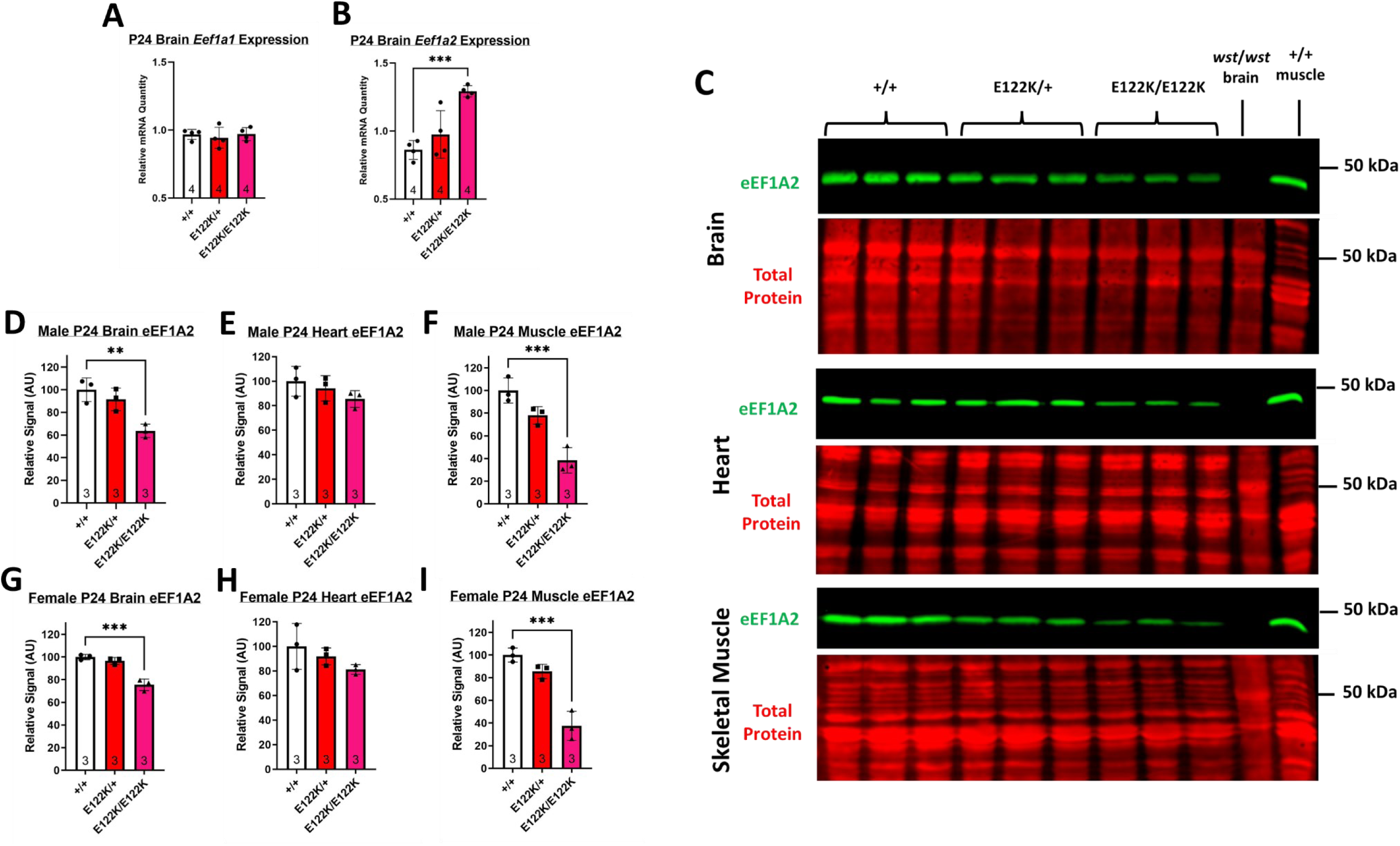
*Eef1a* expression analysis. **A & B**: average *Eef1a1* (A) and *Eef1a2* (B) mRNA quantities in the brains of P24 mice in the E122K line, normalised to the geometric mean of *Gapdh*, *Ubc*, and *B2m* transcript levels. Values for individual biological replicates are shown as dots. Genotypes were compared using ordinary one-way ANOVAs. There was no statistically significant difference in relative *Eef1a1* mRNA quantities between genotypes (F(2, 9) = 0.2807, p = 0.7616). There was a statistically significant difference in relative *Eef1a2* mRNA quantities between genotypes (F(2, 9) = 16.16, p = 0.001). *** denotes p < 0.001 in Dunnett’s post-hoc test (+/+ vs. other genotypes). **C**: representative Western blots of P24 brain, heart and muscle lysates showing eEF1A2 and a total protein stain. Brain lysates from *wst/wst* mice, homozygous for an *Eef1a2* abolishing deletion (Chambers et al., 1998), were used as a negative control for eEF1A2. Lysates from adult skeletal muscle were used as a positive control. **D-I**: quantified eEF1A2 levels, normalised to total protein and expressed as a percentage of wildtype, were compared using ordinary one-way ANOVAs. There were no statistically significant differences in relative eEF1A2 levels in male heart (C; F (2, 6) = 1.595, p = 0.2782) or female heart (F; F (2, 6) = 1.949, p = 0.2227). There were statistically significant differences in relative eEF1A2 levels in male brain (B; F (2, 6) = 13.32, p = 0.0062), female brain (E; F (2, 6) = 39.60, p = 0.0003), male muscle (D; F (2, 6) = 28.49, p = 0.0009) and female muscle (G; F (2, 6) = 39.76, p = 0.0003). ** denotes p < 0.01 and *** denotes p < 0.001 in Dunnett’s post-hoc test (+/+ vs. other genotypes). Sample sizes are shown at the base of the bars. Error bars show the standard deviation.

We next estimated steady state eEF1A2 protein levels in the brain, heart, and skeletal muscle by Western blot (Figure 1C). We found that the E122K mutation reduced average steady-state eEF1A2 levels in all tissues, with significant reduction in E122K/E122K brains and skeletal muscle; however, these reductions were not uniform. Relative to wildtype, eEF1A2 levels in E122K/E122K samples were on average 25-36% lower in the brain, 15-19% lower in the heart, and 62-63% lower in skeletal muscle, with intermediate levels in heterozygotes (Figure 1D-I). The E122K mutation therefore reduces steady state eEF1A2 levels in a tissue dependent manner, likely by decreasing protein stability, with the greatest impact observed in skeletal myocytes. Further Western blotting revealed no significant alteration in eEF1A1 levels in brain, heart, or muscle of E122K/+ or E122K/E122K samples at P24, indicating no compensatory expression/ upregulation of eEF1A1 (Supplementary Figure 4).

### Mice carrying the E122K mutation show growth abnormalities, with homozygotes dying by P31

Analysis of E122K/+ vs. E122K/+ crosses revealed that males and females were born in the expected 1:1 ratio (chi-square test, χ^2^ = 0.25, p = 0.61) and that +/+, E122K/+ and E122K/E122K mice were born in the expected Mendelian ratios in each sex (chi-square tests: male χ^2^ = 1.02, p = 0.60; female χ^2^ = 3.37, p = 0.19). To track early physical development, we measured total body mass of mice in the E122K line between P14 and P28. In order to gain mechanistic insight into the E122K mutation, we simultaneously characterised mice carrying *Eef1a2*-null mutations on the same genetic background (C57BL/6J). These mice were either heterozygous (+/-) or homozygous (-/-) for a 22 bp deletion in *Eef1a2* exon 3 which abolishes expression of eEF1A2 protein (Davies et al., 2020).

While the gross morphology of E122K/+ and E122K/E122K mice was normal, we found that both E122K/+ and E122K/E122K mice developed body mass deficits in early life (Figure 2). E122K/+ growth curves maintained an upwards trajectory but significantly deviated from +/+ from as early as P14 in males and P16 in females (Figure 2A & B); by P28, E122K/+ body mass was on average ∼16% lower than that of wild types in both sexes. Statistically significant body mass deficits in E122K/E122K were detectable as early as P14 in males and P21 in females (Figure 2A & B). E122K/E122K mice gained little weight after P21, and by P28 average E122K/E122K body mass was ∼39% lower than +/+ in males and ∼32% lower than +/+ in females.

**Figure 2.**
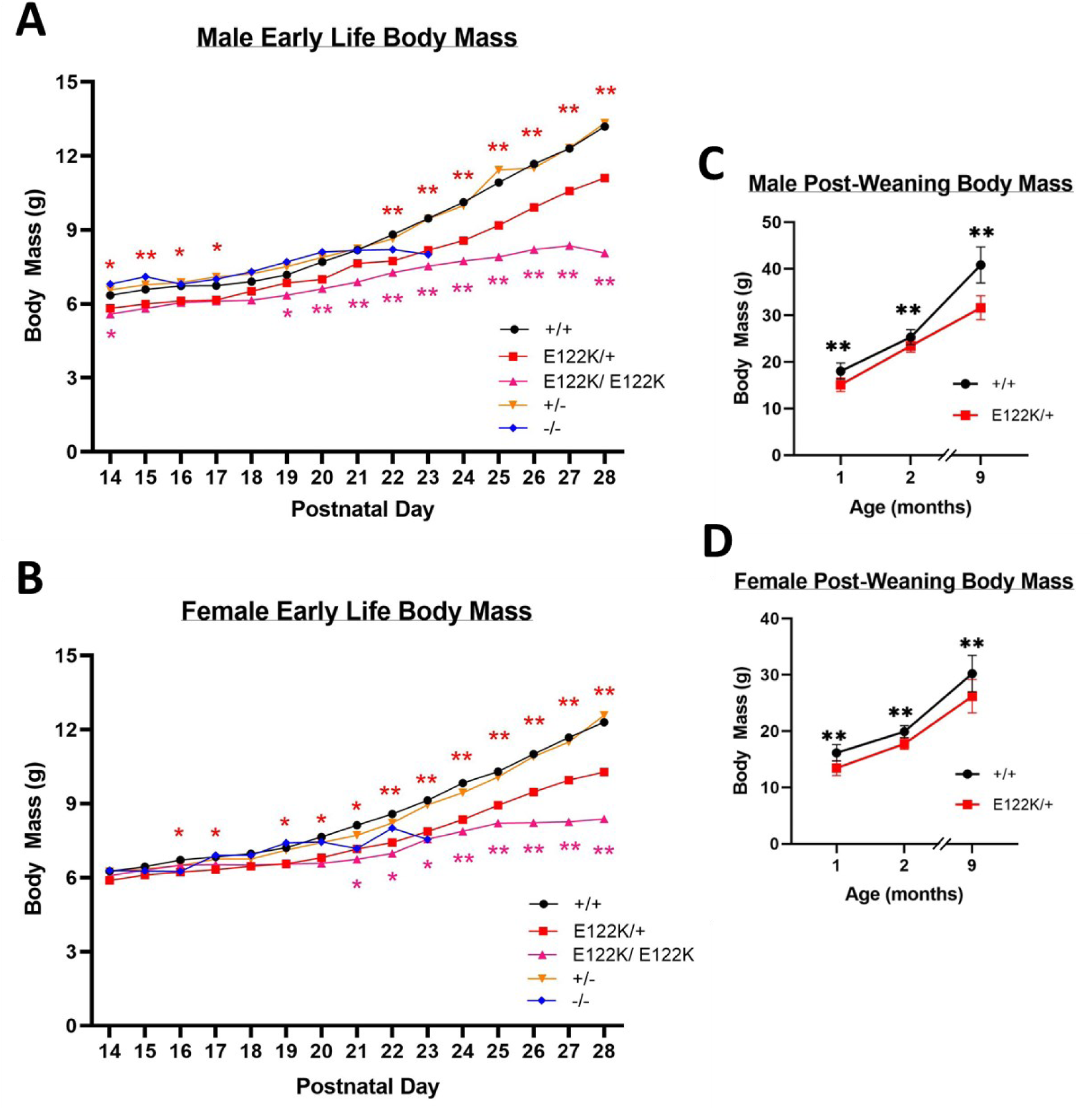
body mass deficits in mice carrying E122K. **A & B**: average absolute body mass in male (A) and female (B) mice between P14 and P28. **C & D**: absolute body mass in male (C) and female (D) mice between 1 and 9 months of age. At each age, genotypes were compared to wildtype using Mann-Whitney U tests followed by Bonferroni’s correction for multiple comparisons. Statistically significant differences are denoted with colour-coded asterisks (* denotes p < 0.05, ** denotes p < 0.01). Error bars in C and D show the standard deviation. Error bars have been omitted in A & B for illustration purposes.

As previously reported (Davies et al., 2020), +/- mice show an essentially identical trajectory to +/+ mice, completely tolerating haploinsufficiency. -/- mice on the C57BL/6J background have stalled weight gain after ∼P20, exhibiting progressive motor/ neurological abnormalities around P18 (Davies et al., 2020; Shultz et al., 1982) and reaching humane endpoints or dying of seizures by P23 (Davies et al., 2017, 2020). The timing of these -/- phenotypes is in line with the postnatal downregulation of eEF1A1 in mice (Chambers et al., 1998; Davies et al., 2020; Khalyfa et al., 2001).

E122K/+ body mass deficits persisted throughout life, with average E122K/+ body mass being ∼23% lower than +/+ in males and 13% lower than +/+ in females by 9 months of age (Figure 2 C & D). Adult E122K/+ mice of both sexes were noticeably leaner than their wildtype littermates, failing to develop the truncal adiposity typically seen in adult laboratory mice and remaining relatively lean even up to 18 months of age, even though average daily food intake did not significantly differ between +/+ and E122K/+ mice at 2 months of age (Supplementary Figure 5). E122K/+ body mass deficits were also not driven by hypotrophy in eEF1A2-expressing tissues, as there was no significant decrease in the relative mass of E122K/+ brains, hearts or *tibialis anterior* muscles relative to body mass (Supplementary Figure 5). The mass of the kidneys (which do not express eEF1A2), were reduced in line with body mass in E122K/+ mice (Supplementary Figure 5), suggesting a consistent and proportional reduction in weight across all tissues.

Complete loss of eEF1A2 function in neurons causes vacuolar degeneration of spinal motor neurons in mice, beginning cervically and progressing rostrocaudally (Newbery et al., 2005) (Supplementary Figure 6C). Surprisingly, we did not observe the same in the cervical spinal cords of E122K/E122K mice at P28, which showed no clear degenerative pathological changes (Supplementary Figure 6A & B). These results indicate that the E122K protein retains sufficient function to spare motor neurons from overt degeneration until at least P28.

### Transient motor delays in E122K/+ mice and progressive motor abnormalities in E122K/E122K mice

To track early motor/ neurological development, we used a composite scoring system called the neuroscore, adapted from (Guyenet et al., 2010) for scoring *Eef1a2* mutant mice (Davies et al., 2020). Cumulative neuroscores were measured between P14 and P60 (Figure 3). In this scoring system, higher scores represent greater motor and neurological dysfunction. In each sex, +/+ mice began with neuroscores of ∼3, diminishing to approximately 0 by three weeks of age as animals physically developed and learned the ledge walking task. The trajectory of +/- neuroscores was essentially indistinguishable from +/+, indicating that haploinsufficiency has no discernible impact on motor/ neurological development. E122K/+ mice of each sex began with similar scores to +/+ at P14 but took longer to decline to a score of zero, remaining marginally but significantly elevated until at least P42 in males and females. The transient elevation of E122K/+ neuroscores was driven primarily by the gait and ledge walking components (Supplementary Figure 7), consistent with early delays in motor development.

**Figure 3.**
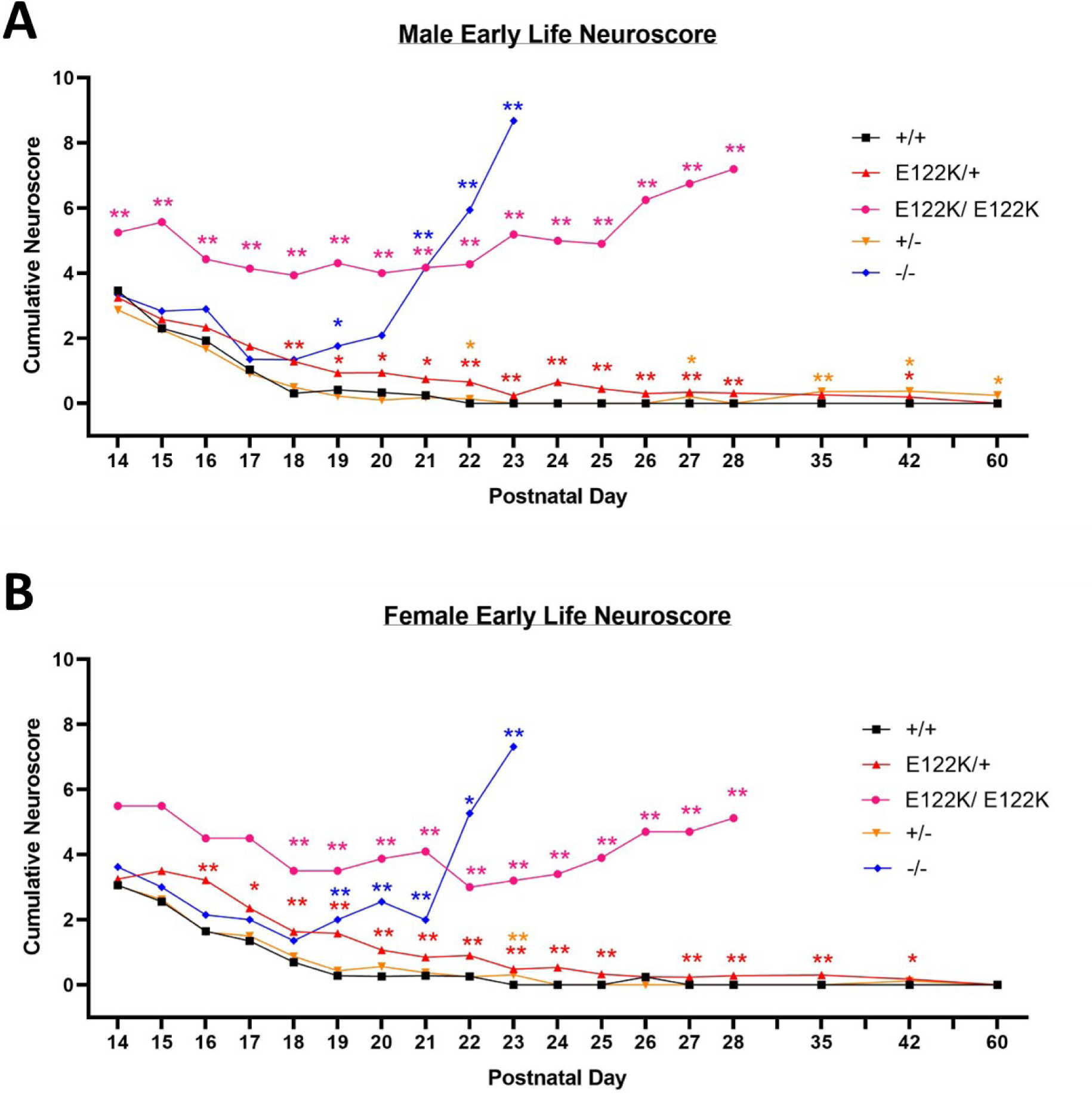
comparative neurological phenotyping reveals that E122K is toxic. **A & B**: average cumulative neuroscores in male mice (A) and female mice (B) between P14 and P60. At each age, genotypes were compared to wildtype using Mann-Whitney U tests followed by Bonferroni’s correction for multiple comparisons. Statistically significant differences are denoted with colour-coded asterisks (* denotes p < 0.05, ** denotes p < 0.01). Error bars have been omitted for illustration purposes.

Neuroscores in null homozygous mice were initially indistinguishable from +/+ but rose precipitously around three weeks of age, in line with the postnatal eEF1A isoform switch in mouse muscle (Chambers et al., 1998; Pan et al., 2004). As previously described, -/- mice develop progressive motor/ neurological abnormalities between 2 and 3 weeks of age, reaching humane endpoints by P23 (Davies et al., 2017). Average E122K/E122K neuroscores were significantly higher than all other genotypes at P14 in males and from P18 in females, remaining significantly higher than those of wild-type littermates thereafter (Figure 4, data shown up to P28). The early E122K/E122K neuroscore was driven by gait abnormalities and difficulty walking along the ledge. Cervical kyphosis emerged around P18, with E122K/E122K mice exhibiting increasingly hunched/ shuffling gaits and progressively losing their ability to walk along a ledge. In addition, E122K/E122K mice developed an essential tremor around P21. In spite of these abnormalities, E122K/E122K mice remained mobile and behaved normally in the cage, but were considered to be at humane endpoints when they progressed from stalled weight gain to weight loss, or began to exhibit reduced movement, declining body condition, grimace, or their tremor began to interfere with movement. E122K/E122K mice reached humane endpoints between P27 and P31.

**Figure 4.**
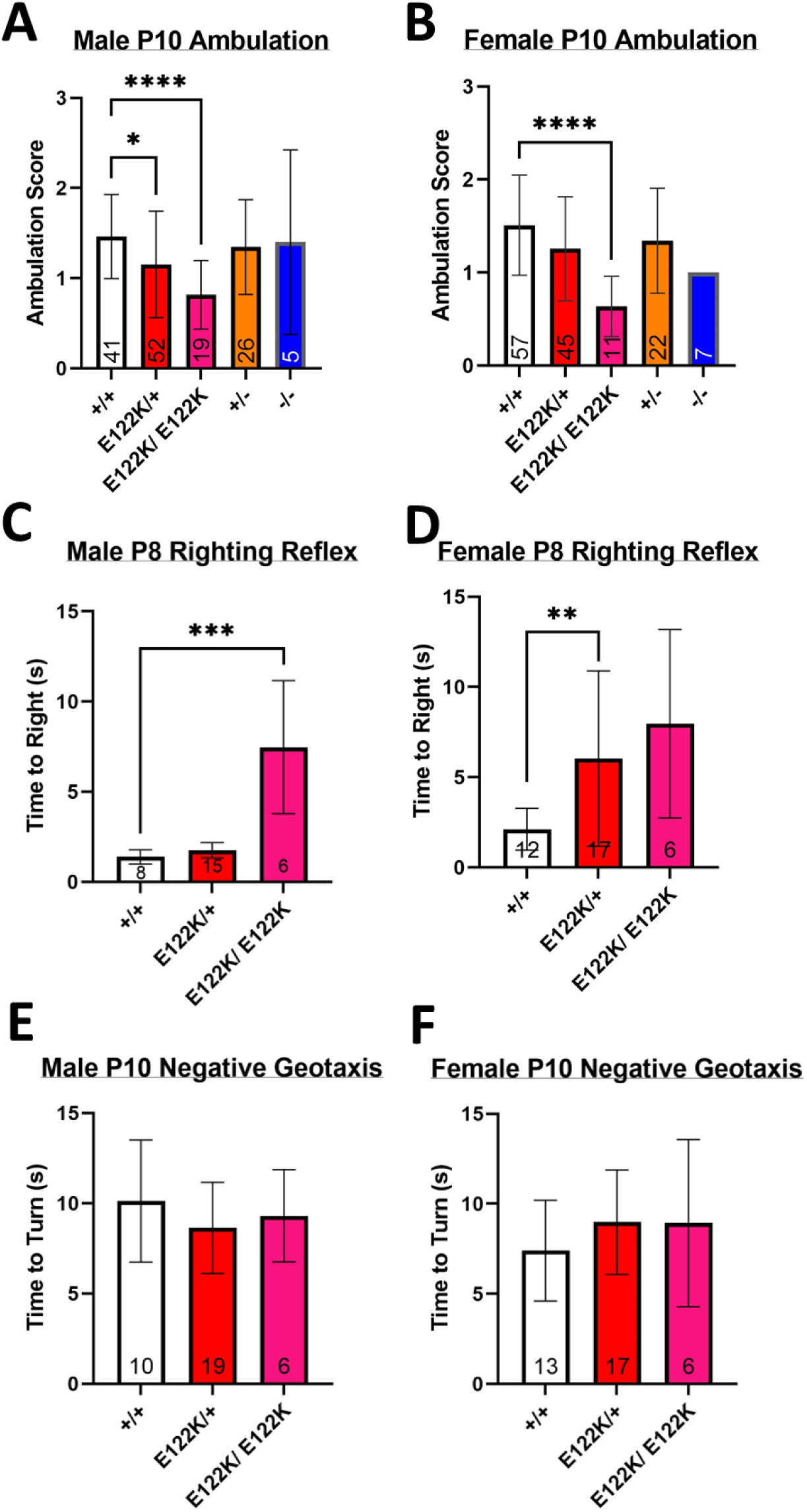
– motor delays in neonates carrying E122K. **A & B**: average ambulation scores in male (A) and female (B) mice at P10. **C & D**: average righting time for male (C) and female (D) mice at P8. **E & F**: average turning time on the negative geotaxis apparatus for male (E) and female (F) mice at P10. Genotypes in A - C were compared using Kruskall-Wallis tests followed by Dunn’s multiple comparisons test as appropriate (+/+ vs. other genotypes, with +/- vs. E122K/+ and -/- vs. E122K/E122K in A & B). There was a statistically significant difference between genotypes in A (KW = 21.7, p = 0.0002), B (KW = 25.57, p < 0.0001) and C (KW = 16.29, p = 0.0003). Genotypes in D were compared using Welch’s ANOVA, revealing a statistically significant difference between genotypes (F(2.000, 11.40) = 4.789, p = 0.0309), followed by Dunnett’s T3 post-hoc test (+/+ vs. other genotypes). Genotypes in E and F were compared using ordinary one-way ANOVAs, revealing no significant differences in males (F(2, 32) = 0.9301, p = 0.4049) or females (F(2, 33) = 1.001, p = 0.3784). * denotes p < 0.05, ** denotes p < 0.01, *** denotes p < 0.001 and **** denotes p < 0.0001 in post-hoc tests. Sample sizes are shown at the base of the bars. Error bars show the standard deviation.

Delayed motor development is a common feature amongst patients heterozygous for E122K (Table 1). To investigate whether the E122K mutation was associated with motor abnormalities in mice, we performed a battery of neonatal motor tests consisting of ambulation scoring at P10, righting reflex tests at P8 and negative geotaxis tests at P10. Test protocols were adapted from Feather-Schussler & Fergusson (2016) (see methods) and were performed by an experimenter blind to genotype. First, ambulation was scored on an ordinal scale by observing P10 neonates as they explored an empty cage. Average ambulation scores for +/+ neonates at P10 were ∼1.5 in both sexes (Figure 4A &B), corresponding to a slow crawl with occasional symmetric limb movement. Ambulation scores in heterozygous and homozygous null mice did not significantly differ from those in wild type mice, consistent with the fact that eEF1A1 remains highly expressed throughout the body at P10. However, ambulation scores were significantly lower in male E122K/+ neonates (mean score of 1.15) and in male and female E122K/E122K neonates (mean scores of 0.82 and 0.64, respectively). These E122K/+ and E122K/E122K scores correspond to [limited] crawling with only asymmetrical limb movement, consistent with a delay in motor development. We next tested the righting reflex in P8 neonates and found that P8 animals of all genotypes exhibited the righting reflex, however both male and female E122K/+ mice took significantly longer than +/+ pups to flip back into the prone position (Figure 4C & D). Wild type neonates of each sex flipped to the prone position in <2.5 seconds on average, while female E122K/+ neonates took on average 6 seconds and male E122K/E122K neonates took on average 7.5 seconds. While these results are not consistent with a delay in reflex development, they suggest that mice carrying the E122K mutation are delayed in developing the strength and/ or coordination to flip from a supine to prone position. Lastly, we performed negative geotaxis tests in P10 neonates. All P10 mice tested were able to detect the incline and turn to face upwards, with no significant differences in the time taken to turn (Figure 4E &F). In summary, this battery of neonatal motor tests has revealed evidence consistent with delayed motor development in mice carrying the E122K mutation, but with no evidence of vestibular deficits.

Hypotonia and progressive motor abnormalities have been found in patients carrying the E122K mutation (Table 1). To investigate whether mice show dystonic phenotypes, we measured grip strength (forelimb and all-limb) between P24 and 9 months of age but found no evidence of consistent grip strength deficits in E122K/+ or E122K/E122K mice (Supplementary Figure 8A-H). To investigate whether E122K/+ mice develop progressive motor abnormalities with age, we assessed their performance on an accelerating rotarod at 2 months of age (as a baseline) and at 9 months of age but found no evidence of declining rotarod performance over this age range (Supplementary Figure 8M-P).

### E122K/+ mice show normal object memory with possible anxiety signatures

To investigate the locomotor activity and anxiety-like behaviours in E122K/+ mice, we performed open field tests at 1, 2 and 9 months of age. We found no statistically significant differences in total distance travelled between +/+ and E122K/+ mice of either sex between 1 and 9 months of age (Supplementary Figure 9A & B), indicating normal locomotor activity levels in out-of-cage contexts. There was no statistically significant difference in thigmotactic behaviour in males (Supplementary Figure 9C). However, female E122K/+ mice showed significantly increased thigmotactic behaviour at 1 and 9 months of age (Supplementary Figure 9D), covering 36% less distance in the centre zone than +/+ mice at 1 month and 45% less distance in the centre zone than +/+ mice at 9 months of age, suggesting increased anxiety levels in E122K/+ females (Gould et al., 2009).

All patients with the E122K mutation exhibit moderate to severe intellectual disability (see Supplementary Table 1). To investigate whether E122K/+ mice exhibited learning/ memory deficits, we administered novel object recognition (NOR) tests at 1, 2 and 9 months of age and determined object discrimination indices for the sample and test trials based on the first 20 seconds worth of object interaction. We found no statistically significant differences in novel object discrimination indices in male or female E122K/+ mice between 1 and 9 months of age (Supplementary Figure 9K-N).

### E122K/+ mice exhibit electrographic seizures and generalised spiking, but do not appear to have convulsive seizures

All patients carrying the E122K mutation have epilepsy, with variable seizure types, ages of onset, and degrees of seizure control (Table 1). Other EEG abnormalities are noted in several patients, including interictal spiking and spectral abnormalities. To investigate whether E122K/+ mice exhibit spontaneous seizures and/ or EEG abnormalities, we performed intracranial EEG recordings in adult mice, with simultaneous EMG recording and video capture.

6 E122K/+ mice (3 male, 3 female) and 6 +/+ mice (3 male, 3 female) were recorded. A total of 124 hours and 36 minutes of EEG data was collected between these 12 animals. The average amount of data collected did not significantly differ between sexes or genotypes (data not shown). Male and female data was combined for all analyses. There were no statistically significant differences between genotypes in the percentage of time spent in wake, NREM or REM, and no statistically significant differences in NREM bout frequency, NREM bout duration or REM bout duration. However, there was a statistically significant 44% decrease in average REM bout frequency in E122K/+ mice (Supplementary Figure 10), suggesting that E122K/+ might have altered REM sleep architecture.

No spontaneous convulsive seizures were observed during EEG recordings of E122K/+ mice, consistent with observations made during behavioural testing. Nevertheless, two putative EEG abnormalities (cortical spike trains and generalised spikes, see Figure 5) were noted during vigilance state classification. The events here termed cortical spike trains (CSTs) were brief and relatively high-amplitude synchronous sharp polyspikes (∼7 Hz spiking frequency) in the motor and somatosensory areas, with occasional generalisation into the parietal cortex (Figure 5B). CSTs were observed in all six E122K/+ mice recorded, with a total of 374 validated events and an average incidence of ∼6 events per hour (95% C.I. 2.9-9.2 events per hour, Figure 5C). CSTs were observed in all vigilance states in E122K/+ mice, although there was significantly lower incidence in NREM sleep (Figure 5C & D). CSTs had a median duration of 0.8 seconds (IQR 0.6-1.1) (Figure 5E), although REM CSTs had a significantly higher median duration than waking CSTs (1.2 seconds (IQR 0.98-1.9) during REM vs. 0.8 seconds (IQR 0.6-1) during waking). A total of 3 cortical polyspike events were observed between two +/+ mice (one male, one female), although these events did match the CST waveform (see Supplementary Figure 11), suggesting that CSTs are physiologically abnormal and possibly ictal events. Indeed, the CST waveform was reminiscent of the spike-wave discharges that are the hallmark of absence seizures (Papale et al., 2009; Pfammatter et al., 2019; Sadleir et al., 2006), yet also strikingly resembled the electrographic seizures observed in *Scn8a* mutant mice (Wagnon et al., 2015). CSTs had no apparent EMG correlates in either wake or sleep and were not consistently associated with altered EMG activity when they occurred during waking, although they often occurred during vigilance state transitions (see Supplementary Figure 11). To investigate whether cortical spike trains had any behavioural correlates, the corresponding video was studied for a random selection of waking CSTs in E122K/+ mice (see Supplementary Video). We found that waking CSTs occurred during a range of normal behaviours, including exploration, rearing, digging, eating, grooming or other stationary activity, and were not associated with convulsions of any sort. The majority of waking CST events investigated coincided with rearing and sniffing, though not all bouts of rearing and sniffing coincided with CSTs. While there appeared to be brief behavioural arrest during some events, behaviour generally continued uninterrupted. As CSTs were not consistently associated with Racine-scoreable behaviour, we conclude that CSTs are purely electrographic seizure events similar to those seen in other mouse models of genetic epilepsy (Wagnon et al., 2015).

**Figure 5:**
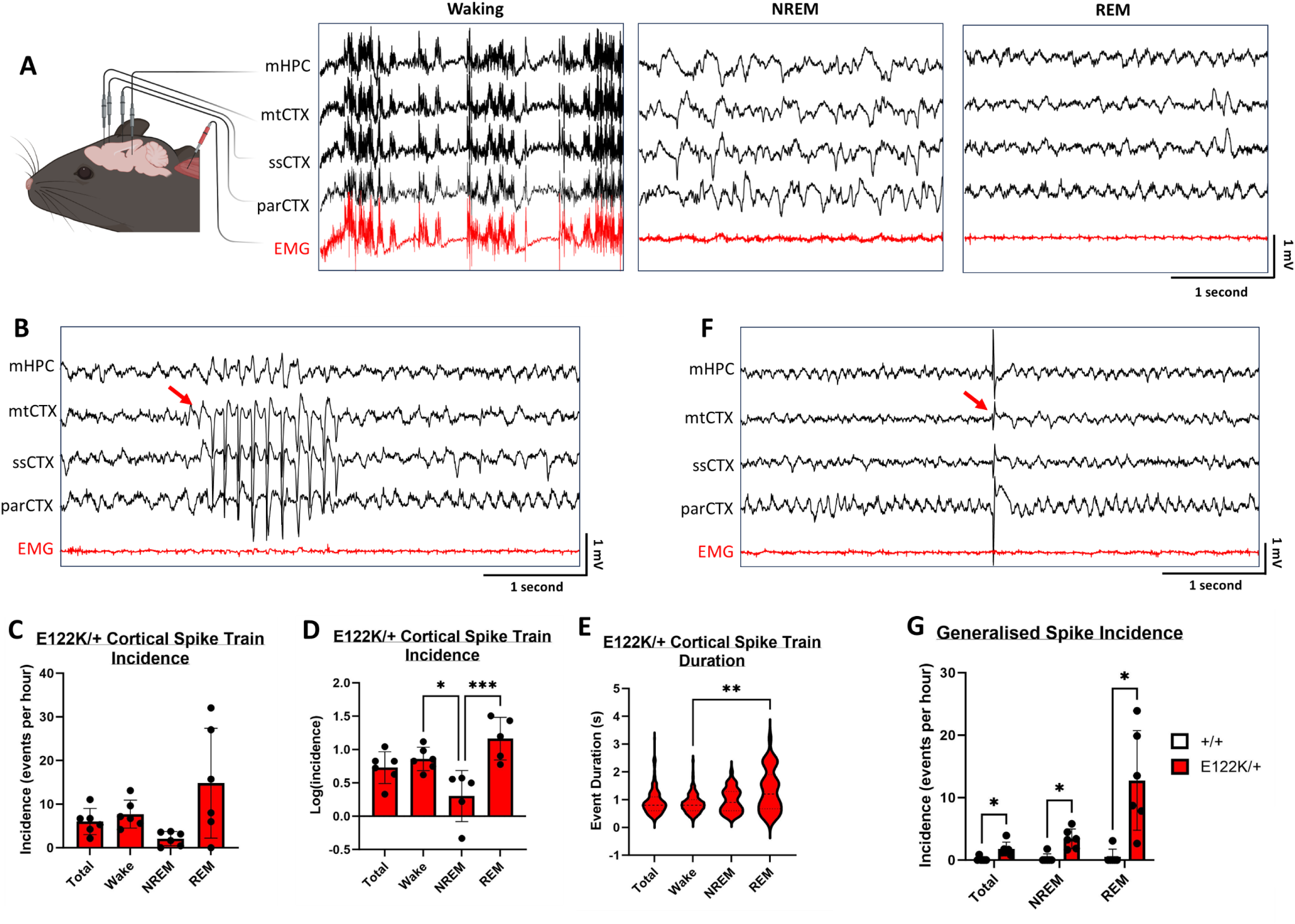
frequent electrographic abnormalities in adult E122K/+ mice. **A**: representative polygraph traces showing EEG and EMG waveforms captured during waking, NREM sleep and REM sleep. Key: mHPC = medial hippocampus, mtCTX = motor cortex, ssCTX = somatosensory cortex, parCTX = parietal cortex, EMG = electromyogram. **B**: 5 second epoch showing a cortical spike train occurring during REM sleep in an E122K/+ mouse. **C**: average incidence (events per hour) of cortical spike trains in total and across each across each vigilance state in E122K/+ mice. **D**: Log transformation of data in C to meet the assumptions of statistical tests. **E**: Violin plots of cortical spike train duration in E122K/+ mice, in total and across each vigilance state. Dashed lines show the median and interquartile ranges. **F**: 5 second epoch showing a generalised spike occurring during REM sleep in an E122K/+ mouse. **G**: average incidence (events per hour) of generalised spikes in total and across each across each vigilance state for +/+ and E122K/+ mice. Vigilance states in D were compared using one-way ANOVA, revealing a statistically significant difference between states (F(3, 18) = 8.040, p = 0.0013). Vigilance states in E were compared using the Kruskall-Wallis test, revealing a statistically significant difference between states (KW = 10.45, p = 0.0151). Genotypes in G were compared within vigilance states using Welch’s t-tests followed by correction for multiple comparisons (Holm-Šídák method). * denotes p < 0.05, ** denotes p < 0.01 and *** denotes p < 0.001. Data in C were not statistically analysed. Error bars show the standard deviation. Panel A mouse illustration created with BioRender.com. Event onsets in B & F are marked by red arrows.

The second EEG abnormality noted during vigilance state analysis was generalised spiking. Generalised spikes were single high amplitude spikes simultaneously affecting all EEG channels but not obviously driven by movement artefacts (Figure 5F). The incidence of generalised spiking was significantly higher in E122K/+ mice (Figure 5G), being observed in all six E122K/+ mice, with a total of 105 validated events and an average incidence of 1.8 events per hour (95% C.I. 0.61-3.0 events per hour). Meanwhile, a total of 8 generalised spikes were observed in a single +/+ mouse (male) giving an average incidence of 0.15 events per hour (95% C.I. -0.22-0.52 events per hour). In contrast to CSTs, generalised spikes were only observed during NREM and REM sleep (Figure 5G). This may be physiologically relevant or simply reflect the fact that waking EEGs are relatively noisy, making it difficult to identify monospikes. Nevertheless, as these events were not identified during waking and did not have obvious EMG correlates, they were also presumed to be entirely electrographic.

Spectral abnormalities have been noted in several patients with the E122K mutation. Notably, increased delta (1-4 Hz) activity in the parietal cortex during the waking interictal EEG has been concomitant with developmental arrest/ regression in some patients ((Inui et al., 2016, 2020), see Supplementary Table 1), representing a possible biomarker of encephalopathy. To investigate whether E122K/+ mice exhibit similar abnormalities, power spectra were generated for each vigilance state and EEG channel, with epochs containing CSTs and generalised spikes excluded. We found no statistically significant differences in power spectral density between genotypes in the delta (1-4 Hz), theta (5-10 Hz), sigma (11-16 Hz), beta (16-30 Hz), low gamma (30-48 Hz) or high gamma (52-100 Hz) spectral bands in any recording location or vigilance state (Supplementary Figures 12 & 13).

## Discussion

Here we used CRISPR/ Cas9 to precisely recapitulate the clinical *EEF1A2* E122K mutation in mice and carried out detailed phenotyping of both homozygotes and heterozygotes, finding that mice carrying E122K exhibit early motor delays, growth defects, and electrographic seizures alongside an increased rate of generalised interictal spiking. To unravel the pathogenesis of DEEs, disease models are required which can reveal the functional consequences of gene mutations. We used direct comparative phenotyping of mice carrying missense or null mutations in *Eef1a2* on the same genetic background, revealing that E122K does not simply phenocopy null mutations. E122K/+ mice exhibit persistent body mass deficits and transient early motor delays that are not seen in +/- mice, and E122K/E122K exhibit earlier onset body mass and neuroscore deficits than -/- mice, indicating that E122K exerts a toxic GOF and/ or dominant-negative effect.

E122K/E122K mice reached humane endpoints between P27-P31, by which point they exhibited neuroscore phenotypes highly reminiscent of that seen in *Eef1a2* nulls. Progressive muscle wastage and motor abnormalities in *Eef1a2* null mice are associated with severe spinal cord pathology which coincides with the downregulation of eEF1A1 (Newbery et al., 2005). Transgenic expression of eEF1A2 in null muscle is insufficient to rescue or even alter the trajectory of the degenerative phenotype (Doig et al., 2013), demonstrating that neuronal LOF is the main driver in the null context. Surprisingly, E122K/E122K mice showed no signs of spinal neurodegeneration by the time they reached humane endpoints, indicating that the E122K protein is not a complete LOF mutation and meaning it is not immediately clear what drives their progressive phenotype. Our Western blots show that the E122K protein is relatively unstable in muscle, with estimated steady state eEF1A2 levels in E122K/E122K mice being ∼60% lower than +/+ in muscle versus only ∼30% lower than +/+ in neurons. Given this and the lack of obvious motor neuron pathology in E122K/E122K mice, we hypothesise that the primary driver of the E122K/E122K phenotype is LOF in muscle.

The molecular bases of any LOF/ GOF have yet to be established. E122K is adjacent to the GTP binding site in the folded protein (see Supplementary Figure 1), meaning it is positioned to affect the kinetics of GTP binding or hydrolysis (Mita et al., 1997; Sandbaken and Culbertson, 1988) and, in turn, the rate of translation elongation. Both increased and decreased translational speeds are associated with protein misfolding, with higher speeds being associated with higher rates of amino acid misincorporation (Wohlgemuth et al., 2010, 2011) and very low speeds associated with ribosome stalling and misfolding of nascent peptides (Ishimura et al., 2014; Komar and Jaenicke, 1995; Nedialkova and Leidel, 2015). Indeed, the E122K mutation was serendipitously characterised in the yeast *EEF1A* orthologue *TEF2*, where it was shown to increase the rate of amino-acid misincorporation (Sandbaken and Culbertson, 1988). Protein misfolding is associated with the accumulation of toxic aggregates (Powers and Balch, 2008), with neurons being particularly vulnerable (Knight et al., 2020). For example, *sticky* mice are homozygous for missense mutations in an alanyl tRNA synthetase causing mischarging of tRNAs, amino acid misincorporation, and protein misfolding, leading to rapid degeneration of cerebellar Purkinje neurons over the first months of life and severe tremors and ataxia (Lee et al., 2006). The E122K mutation also seems to have a neurodegenerative component, with progressive cerebral atrophy and ataxia each noted in patients (see Supplementary Table 1). Nevertheless, there were no obvious signs of neurodegeneration or progressive movement disorders in E122K/+ mice even up to 18 months of age. The toxicity of E122K may also result from disruption of eEF1A2’s non-canonical functions. eEF1A2 has numerous non-canonical roles including modulation of the actin cytoskeleton (Gross and Kinzy, 2005) and co-translational quality control of newly synthesised polypeptides (Gandin et al., 2013). Recently, Mendoza *et al*., showed that the phosphorylation of eEF1A2-specific residues regulates structural plasticity in dendritic spines, tuning actin rearrangements with protein synthesis rates (Mendoza et al., 2021), but there is currently no data on the impact of E122K on any of these processes. One other intriguing possibility is that E122K results in eEF1A2 exerting a dominant-negative against isoform eEF1A1. If this were the case, the effect would be predicted to be most pronounced when both isoforms are co-expressed during development. Interestingly, we observed that the E122K/+ neuroscore gradually declines to insignificance after P21 while the E122K/E122K neuroscore transiently ameliorates around P21, after the eEF1A isoform switch typically concludes (Chambers et al., 1998; Pan et al., 2004). While this might simply reflect the impact of practice, it may also reflect decreasing toxicity as eEF1A1 is downregulated.

The E122K mouse model represents the first animal model of a clinical *EEF1A2* mutation to exhibit relevant, face-valid phenotypes which could be used to assess the efficacy of preclinical treatments. The neonatal motor delays and transient neuroscore abnormalities observed in E122K/+ mice are directly relevant to the delayed motor development and hypotonia observed in patients, while the electrographic EEG abnormalities in adult E122K/+ mice mirror the abnormal interictal spiking observed in patients (see Supplementary Table 1). E122K/+ and E122K/E122K mice also showed robust and persistent growth defects. The E122K/+ mouse model will valuable for preclinical proof-of-concept studies investigating different therapeutic approaches in *EEF1A2*-related neurodevelopmental disorder.

## Acknowledgements

We are grateful to Prof. Colin Smith (The University of Edinburgh) for reviewing the spinal cord sections. The authors gratefully acknowledge the BVS Central Transgenic Core, the staff of the Biomedical Research Facility and the staff of the IGC Advanced Imaging Resource of the University of Edinburgh for their expertise and assistance in this work. We would like to dedicate this paper to the memory of Emma Allan, without whose outstanding skills in microinjection this work could not have been carried out. The work was funded by a grant from the Simons Initiative for the Developing Brain and GFM was supported by a Medical Research Scotland studentship.

